# Arsenic trioxide resistance in acute promyelocytic leukemia: More to it than PML mutations

**DOI:** 10.1101/2020.06.21.154633

**Authors:** Nithya Balasundaram, Saravanan Ganesan, Ezhilarasi Chendamarai, Hamenth Kumar Palani, Arvind Venkatraman, Ansu Abu Alex, Sachin David, Sanjeev Krishna, Anu Korula, Nancy Beryl Janet, Poonkuzhali Balasubramanian, Vikram Mathews

## Abstract

Acquired genetic mutations can confer resistance to arsenic trioxide (ATO) in the treatment of acute promyelocytic leukemia (APL). However, such resistance-conferring mutations are rare and do not explain the majority of disease recurrence seen in the clinic. We have generated a stable ATO resistant promyelocytic cell from a ATO sensitive NB4 cell line. We also noted that another ATRA resistant cell line (UF1) was cross resistant to ATO. We have characterized these resistant cell lines and observed that they significantly differed in their immunophenotype, drug transporter expression, drug resistance mutation profile and were also cross-resistant to other conventional chemotherapeutic agents. The NB4 derived resistant cell line had the classical A216V *PML-B2* domain mutation while the UF1 cell line did not. Gene expression profiling revealed prominent dysregulation of the cellular metabolic pathways in the resistant cell lines. Glycolytic inhibition by 2-DG was efficient and comparable to the standard of care (ATO) in targeting the sensitive APL cell lines and was also effective in the in vivo transplantable APL mouse model; however, it did not affect the ATO resistant cell lines. The survival of the resistant cell lines was significantly affected by compounds targeting the mitochondrial respiration irrespective of the existence of ATO resistance-conferring genetic mutations. Our data demonstrate the addition of mitocans can overcome ATO resistance. We further demonstrated that the combination of ATO and mitocans has the potential in the treatment of non-M3 AML and the translation of this approach in the clinic needs to be explored further.

**Key points:**

- Metabolic rewiring promotes ATO resistance, which can be overcome by targeting mitochondrial oxidative phosphorylation.
- Combination of ATO and mitocans can be exploited as a potential therapeutic option for relapsed APL and in non-M3 AML patients.

## Introduction

Significant advances have been made in the management of leukemia over the last decade. However, resistance to therapy and relapses remains the major obstacle to cure. The major reason for leukemia recurrence has been attributed to the presence of a small population of quiescent leukemic stem cells (LSC), which are inherently resistant to therapy or have acquired genetic mutations that provides a clonal advantage to evade the effects of the therapy ^1-3^. Emerging evidence suggests that additional epigenetic and non-genetic factors in the leukemic cells and their microenvironment, including the immune system, contribute to disease recurrence^4,5^. A better understanding of the mechanisms of resistance has the potential to translate into the development of better therapeutic strategies.

Acute promyelocytic leukemia (APL) is characterized by the presence of reciprocal translocation between the PML gene on chromosome 15 and the retinoic acid receptor α (RARα) gene on chromosome 17 [t(15;17)], resulting in the production of a chimeric and novel PML-RARα fusion oncoprotein that leads to the differentiation block of promyelocytes to mature granulocytes ^6^. In the presence of fusion protein, PML appears to be in a microspeckeled pattern, unlike the wildtype PML where it forms nuclear bodies. PML-RARA degradation restores the p53 signaling and allows the reformation of PML nuclear bodies (NBs). ATO enhances the NBs formation by targeting PML component of the fusion protein, thereby activating the p53 signaling ^7^. p53 and PML nuclear bodies play a critical role in regulating the cellular ROS homeostasis for survival and proliferation^8,9^.

Significant strides made in the understanding of the pathobiology of this disease and the introduction of all-trans retinoic acid (ATRA) and arsenic trioxide (ATO)^10-12^ have contributed to improved survival ^13^. ATO, in combination with ATRA, is considered the standard of care in low and intermediate-risk APL^14^. The current understanding of the mechanisms of action of ATO on the malignant promyelocytes involves ATO induced sumoylation and subsequent poly-ubiquitination of the PML-RARA oncoprotein marking them for proteasomal degradation ^15^. Additional indirect actions include increasing the cellular oxidative damage by ROS, up-regulation of pro-apoptotic factors (JNK, p38 signaling), and downregulating the anti-apoptotic factors (BCL-2, Mcl-1) ^16-19^. Recent observations and studies report novel mechanisms of action of ATO, such as inhibition of glycolysis ^20,21^ and promotion of ETosis (a novel form of cell death) in a dose-dependent manner^22^. These illustrate that the mechanisms of action ATO are complex and multi-factorial. There is evidence supporting the presence of genetic mutations in the *PML* B2 domain (C212-S220; A216V) conferring resistance to ATO in APL, and this is believed to be mediated by altering or inhibiting ATO binding to the B2 domain of the PML component of the PML-RARA oncoprotein^23-25^. However, such acquired somatic mutations are rarely seen in the clinic and cannot explain the majority of relapses that occur in patients treated with ATO based regimens ^26^. In APL, unlike the other subtypes of AML, there is little evidence to suggest the existence of a leukemic stem cell population to explain disease recurrence ^27^.

We have generated an ATO resistant cell line and interrogated its mechanisms of ATO resistance. We report that apart from mutations in the B2 domain of PML-RARA fusion protein, there are non-genetic mechanisms involving cellular metabolic adaptations that contribute to ATO resistance. We further demonstrate that targeting these metabolic adaptations can overcome ATO resistance independent of presence of PML B2 domain mutations.

## Methods

### Cell lines and Chemicals

The human APL cell line NB4 was a kind gift from Dr. Harry Iland, RPAH, Sydney, Australia, with permission from Dr. Michel Lanotte. In addition to our in house generated ATO resistant cell lines, we also used an ATRA resistant APL cell line UF1 (a kind gift from Dr.Christine Chomienne, Hôpital Saint Louis, Paris). The cell lines were free from mycoplasma contamination (Universal Mycoplasma Detection Kit, ATCC Manassas, VA, USA).

Arsenic trioxide kind gift from INTAS Pharmaceuticals, Ahmedabad, India. 2-NBDG – fluorescent analog of D-Glucose, JC-1 and 2-Deoxy Glucose (2-DG), Carbonyl cyanide 4-(trifluoromethoxy) phenylhydrazone -FCCP was purchased from Sigma Aldrich (Sigma Aldrich, St Louis, MO, USA)

### Viability assay

The in vitro cytotoxicity of ATO was determined at 48□h using the MTT assay, as described previously^28^. The viability of the cells was measured post 48 hours of drug treatment using annexin-V-7AAD viability kit (BD Pharmingen, San Diego, CA, USA) as per the manufacturer’s protocol.

### Immunophenotypic Analysis

Immunophenotyping was done on NB4 naïve, ATO resistant NB4 cells, and ATRA resistant UF1 cells. Briefly, cells were labeled using a panel of monoclonal antibodies comprising the myeloid markers directly conjugated with fluorochromes and were incubated for 20 minutes. The cells were then washed and analyzed using FACS Calibur and Cell Quest Pro software (Becton Dickinson, Mansfield, MA, USA).

### Immunofluorescence for PML-RARA protein

Cytospin slides of cell suspensions were made and fixed in 4% paraformaldehyde, followed by blocking using 5% goat serum and incubated with primary antibody (PML PG-M3 -Santa Cruz Biotechnologies, CA, USA) overnight at 4°C. The slides were washed and incubated with a secondary antibody conjugated with Alexa fluor 594 (Invitrogen, Carlsbad, CA, USA) for 1 hour, washed, air-dried, and counterstained with DAPI (4’,6-diamidino-2-phenylindole) containing mountant (Vectashield, Burlingame, CA, USA). The images were acquired in fluorescence microscope (Axioimager M1, Carl Zeiss, Germany) at ×100 with oil immersion and images were analyzed using ISIS metasystem, (Metasystems GmbH, Altlussheim, Germany).

### RNA extraction and qRT-PCR

Total RNA was isolated using TRIzol reagent according to the manufacturer’s protocol (Thermo Fisher Scientific). One microgram of total RNA was used for cDNA synthesis using an iScript cDNA Synthesis Kit (Bio-Rad). Taq-man probe was utilized to quantify the PML-RARA transcripts, and ABL was used as an internal control. The expression of other genes was studied using the SYBR green method (Finnzymes F410L, Thermo Scientific, Rockford, IL, USA). The Ct values were normalized with ACTB, and the fold differences were calculated using the 2^-ΔΔCt^ method.

### Detection of PML-RARA by Immunoblotting

Cell lines were harvested and lysed in RIPA buffer (Sigma) with complete protease inhibitors (Roche, Basel, Switzerland). The lysates were separated through SDS-PAGE and transferred to nitrocellulose membranes (Merck Millipore), blocked in 5% milk in TBS with 0.1% Triton X-100. After blocking the membrane was incubated with primary antibody RARA C-20 (Santa Cruz Biotechnologies, CA, USA) at 4°C overnight, washed, incubated with secondary antibody conjugated with horseradish-peroxidase-anti-rabbit (Cell signaling technologies, MA, USA) for 1 hour, washed and the protein bands were detected by standard chemiluminescence method (Thermo Pierce Femto, Rockford, IL, USA) and the images were captured using FluorChemQ system (Alpha Innotech) provided with Alpha View Software.

### Measurement of Intracellular Arsenic trioxide using atomic absorption spectrometer

The intracellular level of arsenic was measured using well established and published protocols ^29^. Briefly, 2 × 10^7^ cells were washed and suspended in RPMI media with 0.5µM concentration of ATO and incubated for 24 hours. The cell pellets were then washed twice in Ca^2+^ and Mg^2+^ free PBS (Stem Cell Technologies, Vancouver, Canada). The cell pellets were digested with a standard volume of suprapur nitric acid and hydrogen peroxide (Merck Millipore, MA, USA) (2:1 v/v) at 45°C for 16 hours. An aliquot of this final solution was analyzed using Atomic Absorption Spectrometer with EDL support equipment (Perkin Elmer, MA, USA).

### Exome Sequencing

Genomic DNA was isolated from the naïve NB4 cells and the ATO resistant NB4 subclones NB4-EVAsR1 and UF1 using Gentra puregene blood kit (Qiagen, Hilden, Germany) and stored at 4°C. Library preparation and sequencing were performed at Genotypic Technology’s Genomics facility, Bengaluru, following Ion TargetSeq™ Exome Enrichment for the Ion Proton™ System. Sequencing was performed on Ion Proton™ sequencer. The mutations were confirmed by Sanger sequencing.

### Gene Expression array and analysis

A global gene expression array for differential gene expression in naïve NB4 cells and the ATO resistant NB4 primary resistant clone was performed. 2×10^7^ cells (NB4 naïve, NB4 EV-AsR1, and UF1) were harvested and stored in RNA later solution. The extracted labeled RNAs were hybridized to Agilent Human Whole Genome 8×60K Gene Expression Array (AMADID: 039494), and the Image analysis was done using Agilent Feature Extraction software Version 10.5.1.1 to obtain the raw data. Normalization and statistical analysis of the microarray data were done using GeneSpring GX (Agilent Technologies, CA, USA) using the 75th percentile shift and fold difference was calculated by comparing treated samples with control samples. Student t-test and p-values were calculated using the volcano plot algorithm. Differentially regulated genes were clustered using hierarchical clustering to identify significant gene expression patterns Genes were classified based on functions and pathways using biological interpretation tool Biointerpreter (Genotypic Technology; Bangalore). The raw data have been deposited in NCBI’s Gene Expression Omnibus, accessible through GEO Series accession number GSE115812 (https://www.ncbi.nlm.nih.gov/geo/).

### Endogenous Reactive Oxygen Species (ROS) measurement

Endogenous ROS levels of the cell lines were measured by labeling 5×10^5^ cells for 30minutes at 37°C with redox-sensitive probes CellRox (5µM) (Life Technologies, NY, USA). Stained cells were washed twice with PBS and analyzed using Beckman Coulter Gallios flow cytometer (Becton Dickinson, Mansfield, MA, USA, and Kaluza Analysis Software (Becton Dickinson, Mansfield, MA, USA).

### Measurement of Mitochondrial Membrane Potential (MMP)

The MMP (ΔΨm) of the cells was measured using JC-1 dye (Life Technologies, Carlsbad, CA, USA), as previously reported. The fluorescence intensity was measured using Spectramax M4 (Molecular Devices, Sunnyvale, CA, USA) (green channel: excitation: 485□nm; emission: 530□nm; cutoff 515□nm; red channel: excitation: 485□nm; emission: 590□nm; cutoff 570□nm).

### Seahorse Extracellular Flux analyzer

Extracellular flux assay kits XF24 (Agilent Technologies, CA, USA) was used to measure oxygen consumption and glycolytic flux. Briefly, three replicate wells of 5□×□10^4^ cells per well were seeded in a retronectin (Takara Bio Inc, JPY) coated 24-well XF24 plate. At 30□min before analysis, the medium was replaced with Seahorse XF media (Agilent Technologies, CA, USA), and the plate was incubated at 37□°C. Analyses were performed both at basal conditions and after injection of glucose, oligomycin, and 2-deoxy glucose for glycolytic function. Extracellular acidification rate (ECAR) indicates glycolytic capacity. Seahorse extracellular flux analysis was carried out at the National Centre for Biological Sciences (NCBS, Bengaluru) and Department of Biotechnology, Anna University, Chennai.

### Mouse model and drug treatments

FVB/N mice were obtained from Jackson Laboratory (Bar Harbor, ME, USA). Mice at 6 to 8 weeks of age were used in all the experiments. The animal study design and euthanasia protocols were approved by the institutional animal ethics committee (IAEC approval number 04/2019). APL cells from the spleen of MRP8-PML-RAR transgenic mice (FVB/N) were harvested and cryopreserved (a kind gift from Dr. Scott Kogan, UCSF, USA). APL cells (10^6^ cells/mouse) were injected intravenously via the tail vein into genetically compatible FVB/N recipients, without conditioning with either radiation or chemotherapy. After the leukemic cell engraftment period (day 8), intraperitoneal injection of ATO (10mg/Kg) and 2-DG (750mg/kg) was given for 15 days.

### Statistical Analysis

All data points are represented as means ± SEM. Two-tailed Student’s t-test was used to compare mean values between two groups. One-way or two-way analysis of variance (ANOVA) was used to compare mean values between multiple groups. Statistical analysis was performed using Prism version 7.0 (GraphPad Software, La Jolla, CA). P values < 0.05 were considered statistically significant.

## Results

### Generation of arsenic trioxide resistance cell line

ATO resistant cells were generated by exposing the naïve NB4 cell line to low concentrations of ATO (50nM) for about 3 months. Once the cells persisted and sustained the concentration of ATO was gradually increased to 1uM over one year. The cells which survived and proliferated at 1uM were termed as the “ATO tolerant persister cells” (ATO-TPs). The ATO-TPs were then subjected serially to limiting dilutions and single cell colony-forming unit formation on methylcellulose to isolate monoclonal resistant populations. We isolated three different clones, expanded and named them NB4 EV-AsR1, NB4 EV-AsR2, NB4 EV-AsR3 respectively based on the published norms of NB4 resistant cell line nomenclature^30^. The IC50 for ATO in these cell lines was 3.25uM, 3.4 μM, and 2.88 μM for NB4 EV-AsR1, NB4 EV-AsR2, and NB4 EV-AsR3 respectively in contrast to naïve NB4 which was 0.9uM (figure 1a). The viability of the in-house generated ATO resistant cell lines were not significantly affected by exposure to 2uM ATO in comparison to the sensitive cell line NB4 (figure 1b). We have also observed that UF1, a known ATRA resistant cell line, was cross resistant to ATO with an IC-50 of 4.9uM, and this observation has not been previously reported (Supplementary figure 1). The in-house generated ATO resistant cell lines, and UF1 were also significantly less sensitive to differentiation-inducing agent ATRA (figure 1c).

**Figure 1:**
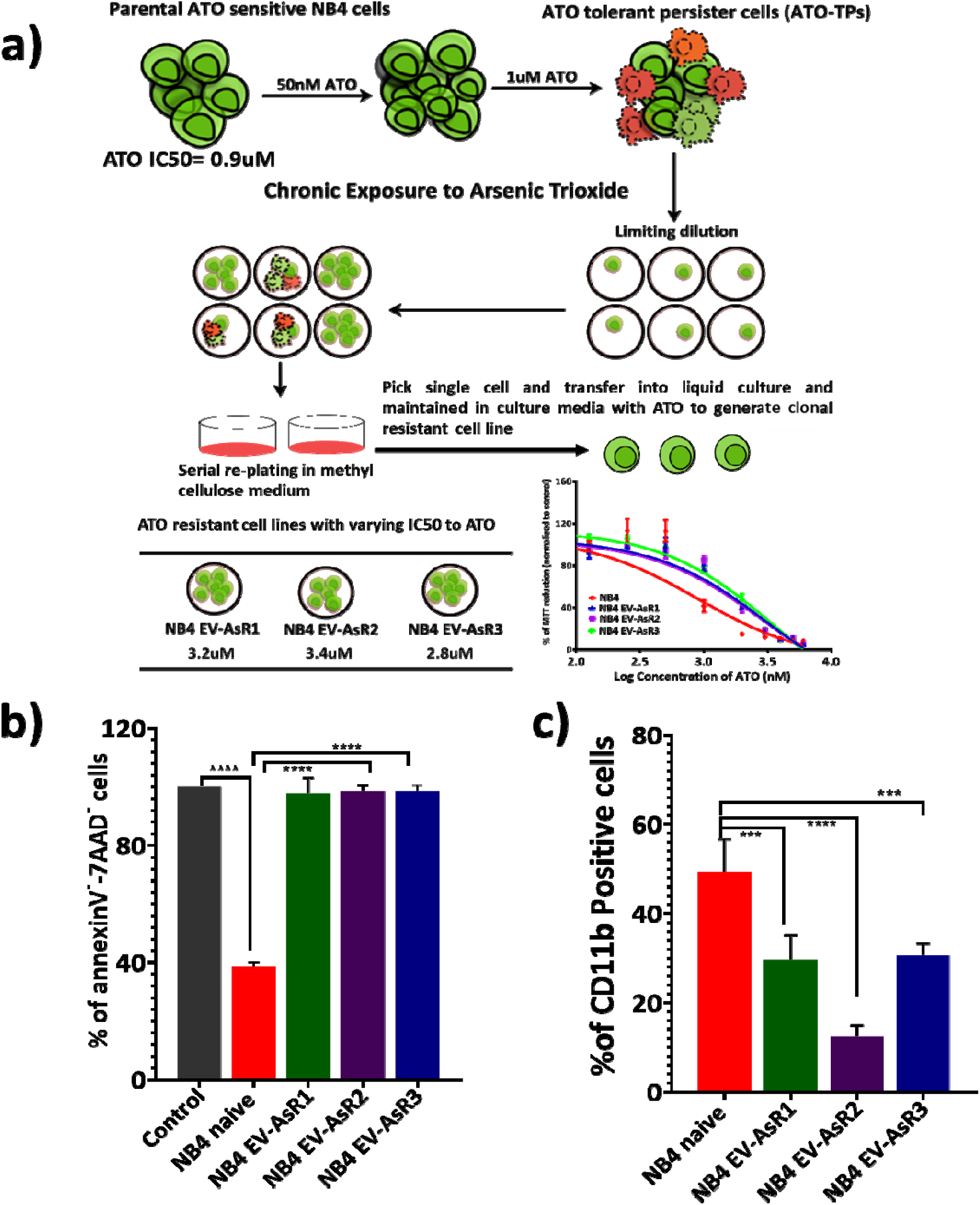
Generation of ATO resistant cell lines. a) NB4 naive parental cell line was exposed to 50nM of ATO for three months, and the concentration was gradually increased to 1uM ATO over a period of one year until they sustained and proliferated. Limiting dilutions and colony-forming unit assay were performed to generate mono clones of the resistant cell lines. b) The bar graph represents the percentage of viable cells post 48 hours of 2uM ATO. c) NB4 and resistant cell lines were treated with 0.5uM of ATRA for 72hrs, and the percentage of differentiation was measured by the surface expression of CD11b. Graphs and statistical parameters were generated from four independent experiments. *p≤0.05; **p≤0.01; ***p≤0.001; ***p≤0.0001.

The doubling time of parental cell line NB4 was 28 hours, whereas, for NB4 EV-AsR1 and NB4 EV-AsR2 clones, it was 46 and 48 hours, respectively. Long term withdrawal of ATO for 3 months from the culture system did not result in reacquisition of ATO sensitivity, and the inhibitory concentration did not change after prolonged and repeated drug holiday (supplementary figure 2). Currently, the resistant cell lines have a stable resistant phenotype when grown with or without ATO.

**Figure 2:**
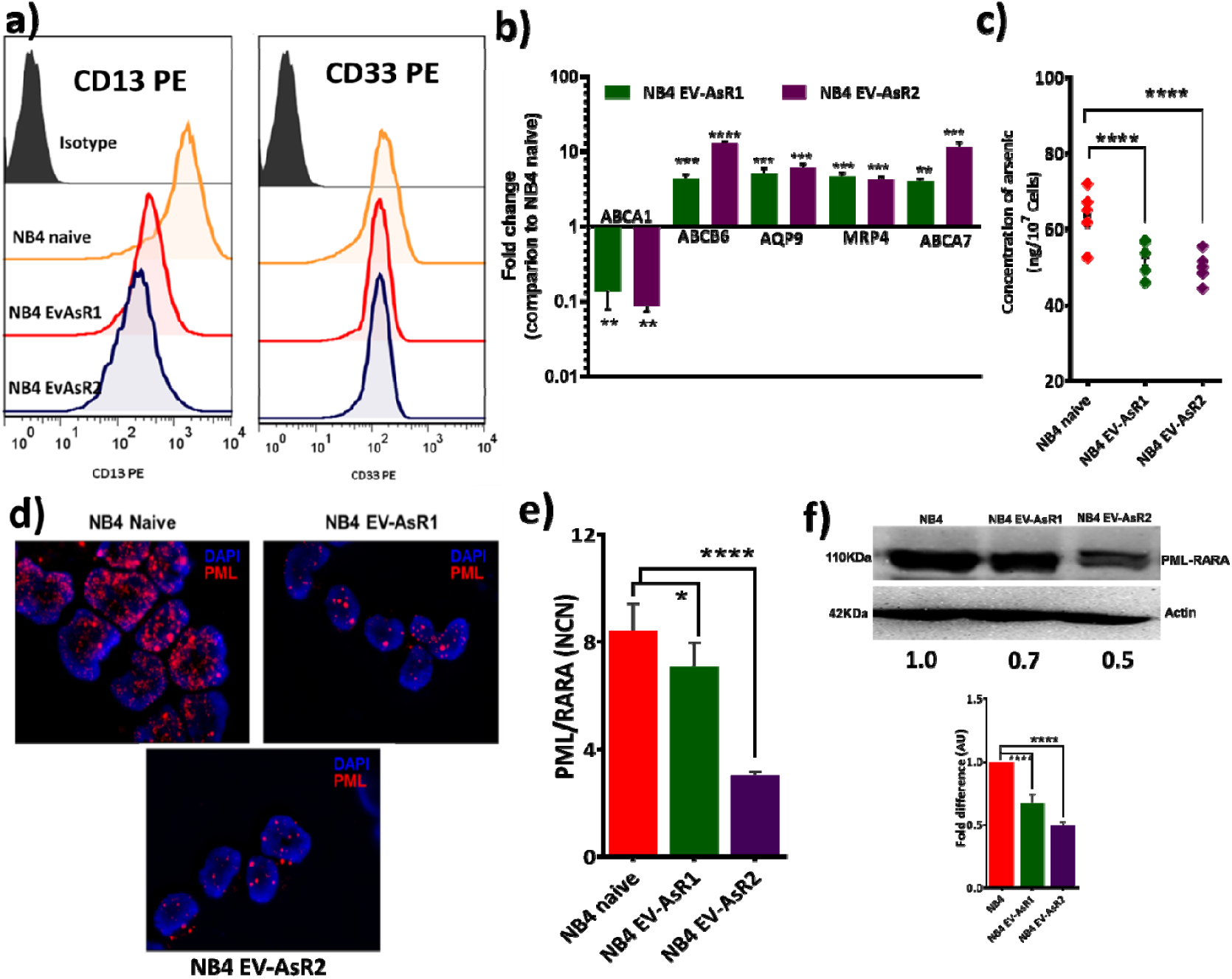
Heterogeneity in the cell surface marker and transporters expression of the ATO resistant cell line in comparison to parental cell line: a) Representative histograms of cell surface expression of CD33, CD13 were assessed by flow cytometry (n=3). b) Relative mRNA levels of ATO influx gene (ABCA1) and efflux transporters genes (ABCA7, AQP9, MRP4, and ABCB6) in in-house generated ATO resistant cell lines compared to NB4 naïve which is normalized to one (n=3). c) Intracellular ATO levels in NB4 naïve and ATO resistant cell lines post 24 hours of 0.5uM ATO treatment (n=4). d)Fluorescent microscopic image of the PML (RED) in NB4 naive and ATO resistant cell lines displaying nuclear body formation and microspeckeled pattern (magnification 63x – oil immersion) e) PML-RARA transcripts in the NB4 naive and inhouse generated ATO resistant cell lines (n=6) f) Immunoblots of the fusion protein levels and fold change in comparison to NB4 naive. All error bars represent the mean ± SEM of three independent experiments. *p≤0.05; **p≤0.01; ***p≤0.001; ***p≤0.0001.

### ATO-resistant cell lines exhibit distinct cell surface markers and transporters

The ATO resistant cell lines were abnormal promyelocyte where the cell surface expression of the typical myeloid markers CD13 and CD33 was significantly reduced in comparison to the parental naïve cell line (figure 2a). The ATO efflux transporters such as AQP9, MRP4, and MRP9 have significantly up regulated in the ATO resistant cell lines which mirrored in the reduced concentration of ATO intracellularly (figure 2b and 2c).

On PML immunofluorescence studies, we noted nuclear body (NB) formation in the resistant cell lines, unlike the naive NB4 cells where these were completely absent, and the characteristic micro-speckled pattern was seen (figure 2d). The PML-RARA transcripts and protein levels were also found to be significantly lower in the ATO-resistant cell lines in comparison to the NB4 cell line (figure 2e and 2f).

### In-house generated ATO resistant cell lines acquired additional cytogenetic and molecular aberrations

In addition to the *PML-RARA* chromosomal translocation, NB4 EV-AsR1 acquired additional cytogenetic abnormalities such as a second small hyper diploid clone (chromosome number 56∼62), deletion 5q, gain of chromosome 4, loss of chromosome 22 and no loss of chromosome X, absence of the derivative chromosome 21, absence of addition 16q and presence of +3 and +12 in the hyper diploid clone were found to be unique in the ATO resistant cell line (supplementary table 1)

As there are reports implicating the emergence/existence of drug resistance-conferring (PML domain mutations) somatic mutations in APL cells against ATO, we performed whole-exome sequencing on our in-house generated ATO resistant cell line NB4 EV-AsR1(as a representative of other ATO resistant clones) in comparison to the parental cell line NB4 naïve and also on the UF1 (ATRA resistant cell line) found to be resistant to ATO.

Whole exome sequencing revealed that in comparison to NB4 naive, a significant number of genes were mutated in the NB4 EV-AsR1 and UF1 cell lines. Based on the mutation frequency, we observed that in comparison to the NB4 naïve majority of the mutated genes in NB4 EV-AsR1 belong to cell surface proteins, especially mucins (MUC6, MUC5B, MUC4, MUC3A, MUC16) and PRSS genes (PRSS1, PRSS3, PRSS3P2). While the UF1 cell line showed a higher frequency of mutations involving MUC16, ITGB4, PRSS1, CUL7, CDH23, LTBP3, OBSCN, STAB1 genes (supplementary figure 3).

**Figure 3:**
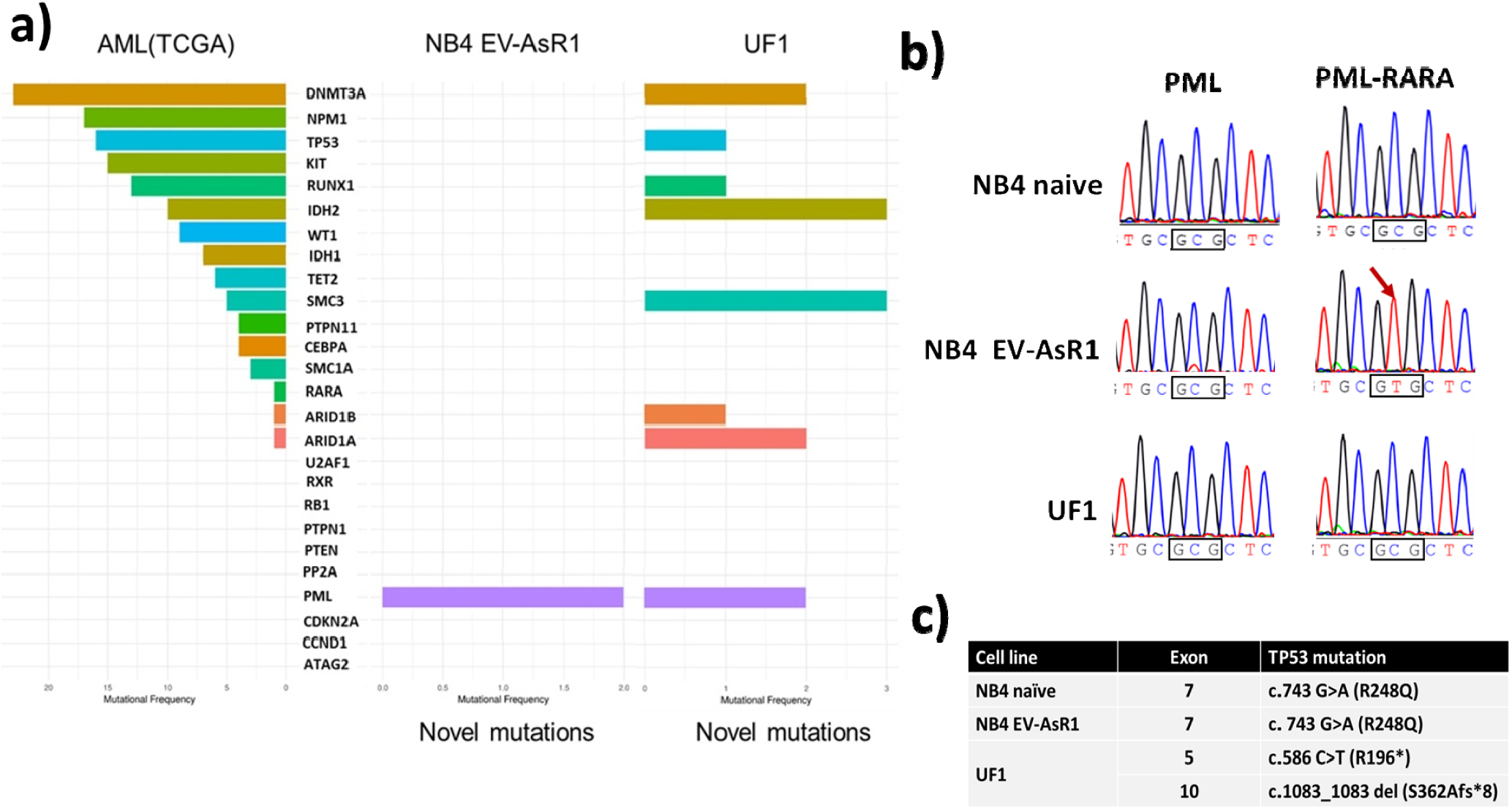
Whole-exome sequencing reveals changes in the ATO resistant cell lines at the genomic level. a) AML TCGA data set was compared with novel mutations observed in of AsR1 and UF1, and the graph represents the novel mutations (only found in resistant cell lines) and their mutation frequency. B) Sanger sequencing showed the existence of PML – A216V in the inhouse generated ATO resistant cell line and not in the UF1 and parental cell line NB4 naïve. c) Table represents the mutation observed in the p53 gene.

While focusing on ATO resistance, we observed the presence of PML B2 domain ATO resistance-conferring mutation A216V in the NB4 EV-AsR1 and not in the UF1 cell line. Further comparison of mutated genes in the resistant cell lines with the commonly observed mutations in the AML TCGA dataset revealed that the resistant cell line had gained additional and novel mutations in them over and above those seen in the naïve NB4 cell line. In NB4 EV-AsR1, except the PML, we did not observe any additional mutation in the TCGA gene set. However, in UF1, there were additional novel mutations observed in the DNMT3A, TP53, RUNX1, IDH2, SMC3, ARID1B, ARID1A, and PML genes addition to the known mutations of AML TCGA gene set (figure 3a).

We validated the PML and p53 mutation using Sanger sequencing, where the ATO resistance-conferring mutation A216V was present in the in-house generated ATO resistant cell lines. UF1 had two intronic variations in the PML domain and was negative for A216V (figure 3b). We also noted that the existence of a p53 gain of function mutation (R248Q) in the in house generated ATO resistant cell line, which was also present in the parental naïve NB4 cell line. UF1 cell line had a point mutation (R196*) and a deletion of exon 10 of the p53 (figure 3c), which are reported to be pathogenic.

### Non-genetic heterogeneity and ATO resistance

As it was evident from drug withdrawal conditions and exome sequencing analysis, the observed ATO resistance was not due to a transient epigenetic poising or known acquired genetic mutations in the *PML-B2* domain (absence of *PML-B2* domain mutation in UF1). We subjected NB4 naïve, UF1, and one of the ATO resistant sub-clone (NB4 EV-AsR1) to gene expression profiling. We observed that 1717 genes in NB4 EV-AsR1 and 6149 genes in UF1 genes were significantly upregulated (> 2-fold) in comparison to NB4 naïve. The pathways significantly enriched for differentially expressed genes were cell survival, cell cycle, immune regulation, ABC transporters, glutathione metabolism, redox system, mitochondrial biogenesis, cellular respiration, and ubiquitin-proteasome degradation system. We also noted that the gene expression profile of the in-house generated ATO resistant cell line was similar to the gene expression profile of relapsed APL patients treated with front line ATO based regimens (Figure 4b).

**Figure 4:**
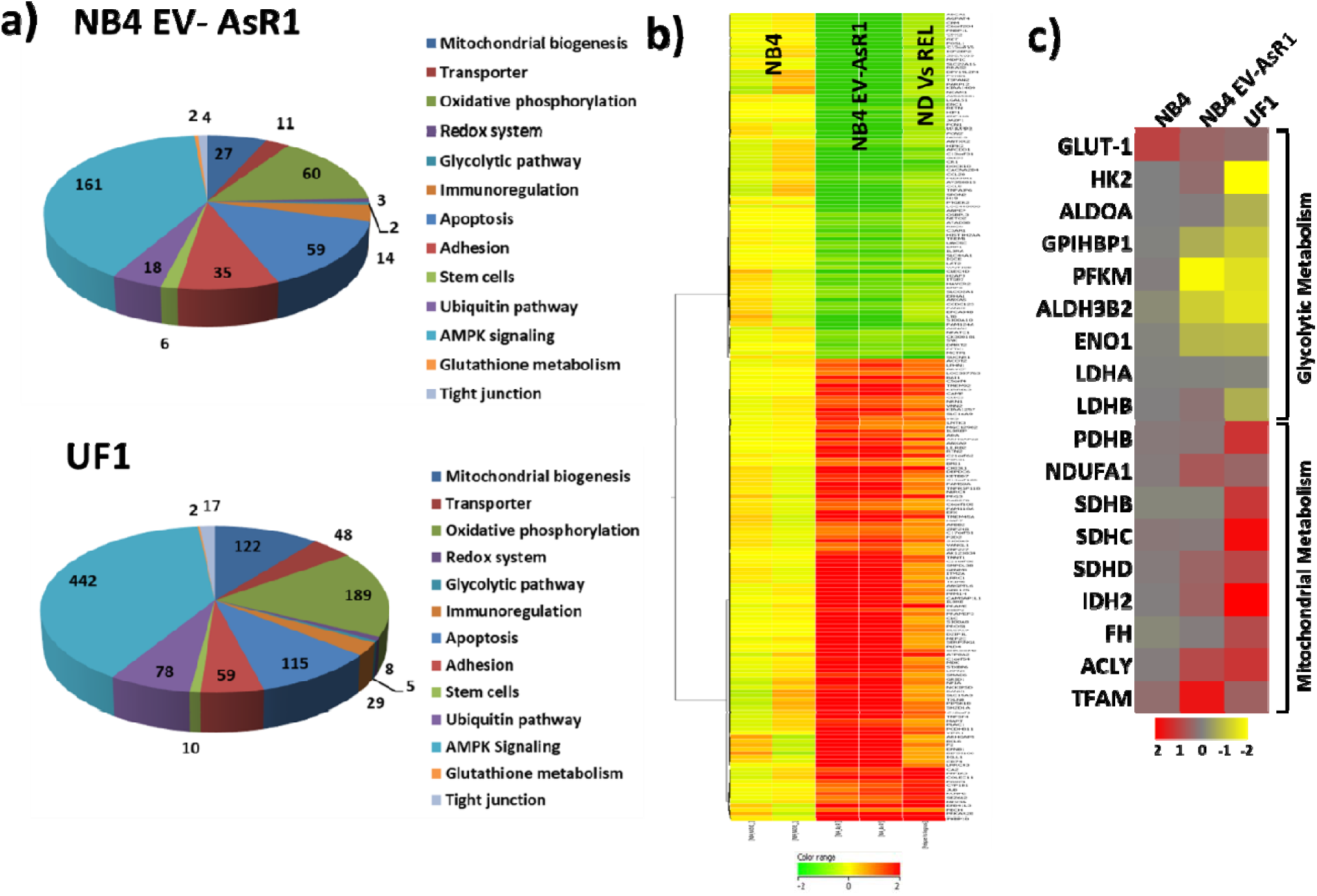
Gene expression analysis of the ATO resistant cell lines reveals dysregulation of cellular metabolism. a) Pie chart representing the dysregulated genes in the inhouse generated ATO resistant cell line NB4 EV-AsR1 and UF1 in comparison to parental cell line NB4 naive. b) Heatmap showing commonly dysregulated genes between ATO resistant cell lines and relapsed APL samples. c) Heatmap representing the genes involved in the glycolytic and mitochondrial metabolism genes.

Gene expression profiling revealed significant dysregulation of glycolytic and mitochondrial metabolism in the resistant cell line when compared to NB4 naïve (figure 4c). We validated the finding by measuring the basal metabolic properties such as reactive oxygen species (ROS), antioxidant level, glucose uptake, and mitochondrial membrane potential (MMP), which are also reported to be key factors in the mechanism of action of ATO. We observed that in comparison to naïve NB4 cells, ATO resistant cell lines had decreased levels of ROS and increased levels of glutathione (GSH) (Figure 5a and b). Mitochondrial membrane potential (MMP) and glucose uptake capacity of the resistant cell lines were observed to be significantly lower in comparison to the naïve NB4 cells (figure 5c, d, and e).

**Figure 5:**
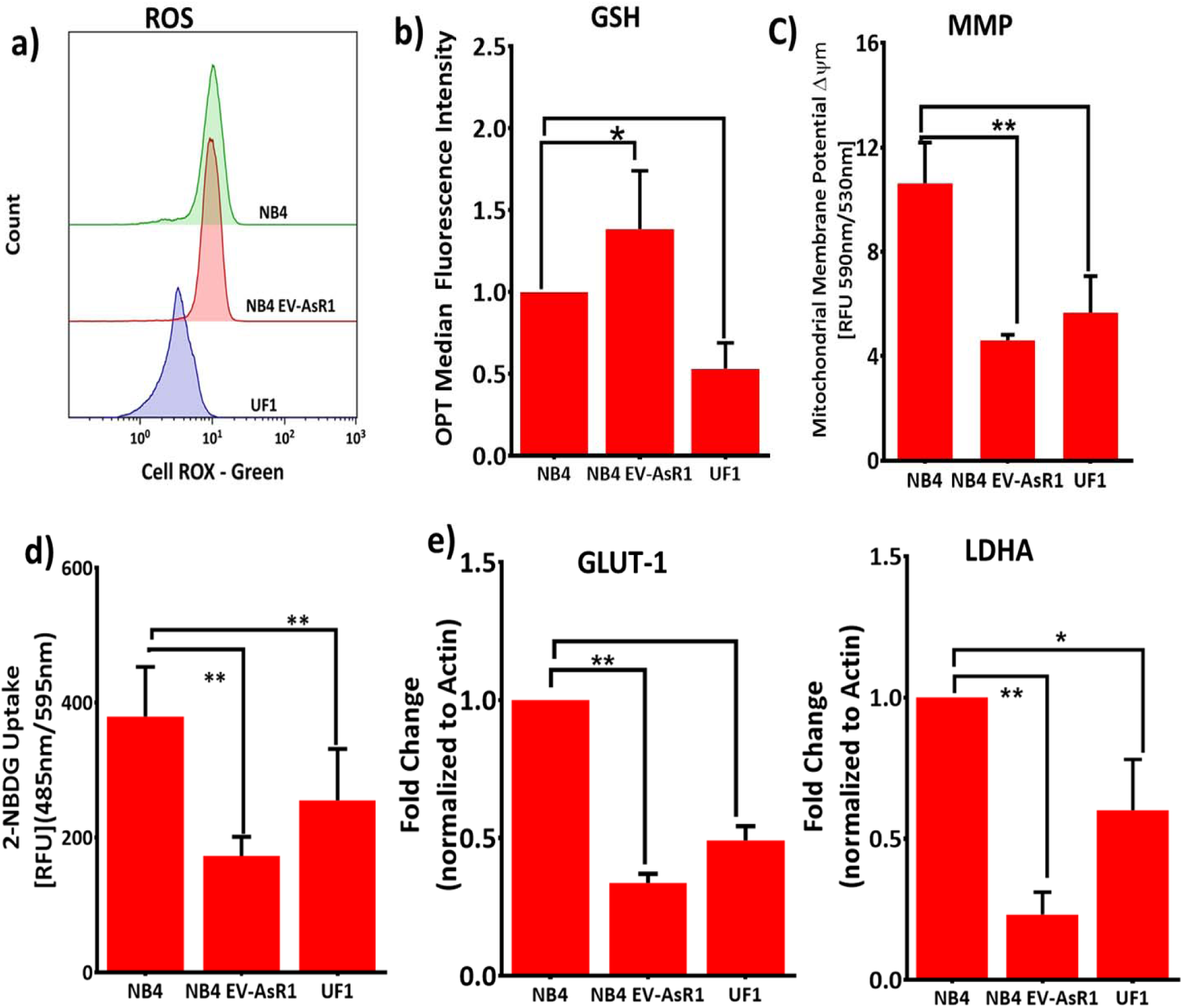
ATO resistant cell lines are metabolically distinct. a) Baseline total reactive oxygen species were measured using redox-sensitive dye – cell ROX Green in the flow cytometry (n=3). b) Baseline protein thiols was measured as an indicative of antioxidant using OPT (Phthaldialdehyde) and median fluorescence intensity are represented as bar graphs(n=3). c) Mitochondria membrane potential of the resistant were measured using JC-1 d) Glucose uptake was measured using a fluorescent analogue of 2-deoxy glucose and represented as relative mean fluorescence intensity (n=3). e) GLUT-1 and LDHA transcripts were signicantly less in the resistant cell lines (n=5). All error bars represent the means ± SEM of three independent experiments. *p≤0.05; **p≤0.01; ***p≤0.001; ***p≤0.0001.

### ATO resistant promyelocytic cells are metabolically distinct irrespective of PML mutation A216V

Seahorse analysis found that high MMP naïve NB4 cells had an increased glycolytic rate (ECAR: extracellular cellular acidification rate was increased as a result of increased lactate production) in comparison to the MMP low ATO resistant cell lines (figure 6a).

**Figure 6:**
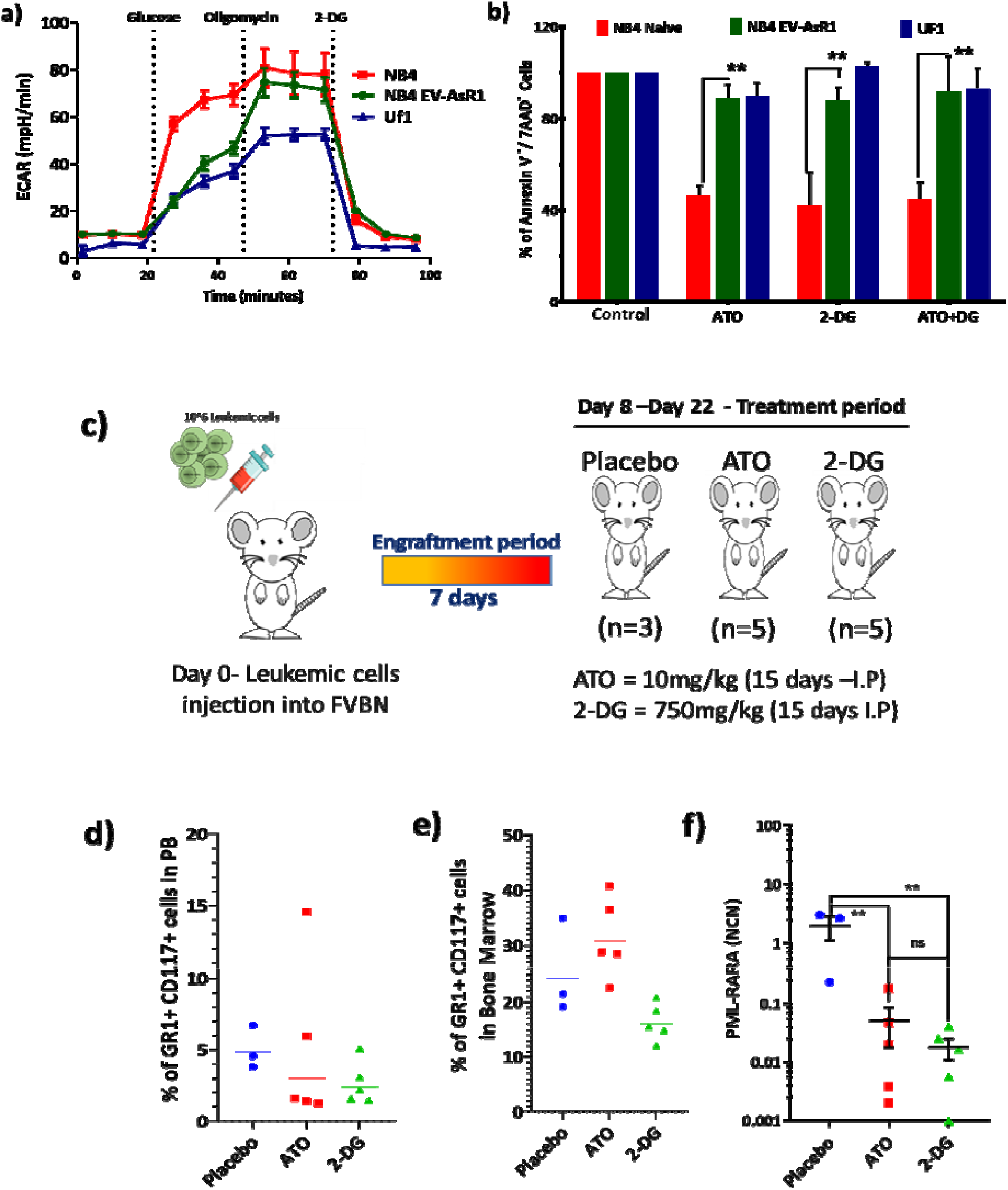
ATO resistant cells are metabolically heterogenous and *in vivo* effect of glycolytic inhibition by 2-DG reduces leukemic burden in APL mouse model. a) Extracellular acidification rate (ECAR) in NB4 naïve, NB4 EV-AsR1 and UF1 cell line. b) Viability of the sensitive and resistant cell lines post 48 hours of glycolytic inhibitor ATO and 2-DG (n=4). c) Schematic representation of the APL transplantable mouse model and treatment plan. Mice were euthanized on day 22 and examined for the presence of leukemic cells as CD117+Gr1+ cell in d) peripheral blood, e) bone marrow and f) PML-RARA transcripts levels in bone marrow.

To address the degree to which glycolysis is necessary, we treated the naïve NB4 and ATO resistant cell lines with 2-Deoxy glucose (2-DG), a glucose analog that inhibits glycolysis via its action on hexokinases. We noted that naïve NB4 cell line viability was significantly affected in the presence of a glycolytic inhibitor 2-Deoxy glucose (2-DG) equivalent to the effect seen with 2uM of ATO and there was no evidence of an additive effect when these agents were combined (figure 6b). In contrast, the viability of the resistant cell line was not significantly affected when the 2-DG was used alone at the same concentrations or when it was combined with ATO (figure 6b). The data suggest that the ATO resistant cell lines in contrast to ATO sensitive cell lines were not relying on the Warburg effect, for their proliferation and survival.

Having noted that glycolytic inhibition by 2-DG promoted apoptosis in NB4 cells comparable to that seen with ATO, we performed in-vivo glycolytic inhibition to understand the physiological relevance of glycolytic inhibition in the ATO sensitive transplantable APL mouse model (figure 6c). We observed that 2-DG or ATO as single agents reduced the leukemic burden in the peripheral blood (PB)(figure 6d), bone marrow (BM)(figure 6e), and PML-RARA copy number in BM (figure 6f) at the end of 22 days to levels that were comparable and indistinguishable from each other.

### Combination of glycolytic inhibitor (2-DG / ATO) and mitocans promoted apoptosis in the ATO resistant cell lines

We then evaluated the effect of mitochondrial OXPHOS inhibitor and glutathione synthesis inhibitor (buthionine sulfoxamine; BSO) on the ATO resistant cell lines. The viability of ATO resistant cell lines was not affected significantly when treated with the mitochondrial OXPHOS un-coupler FCCP or BSO as single agents, whereas in combination with ATO, the viability was significantly reduced in both the A216V mutated NB4 EV-AsR1 and A216V negative cell line UF1(Figure 7a). We observed that the ATO resistant cells lines were significantly different in their metabolic preferences for their survival from the ATO sensitive NB4 naïve cell line (figure 7b) and targeting this difference was able to overcome the resistance independent of the presence or absence of the *PML-B2* domain mutations associated with ATO resistance

**Figure 7:**
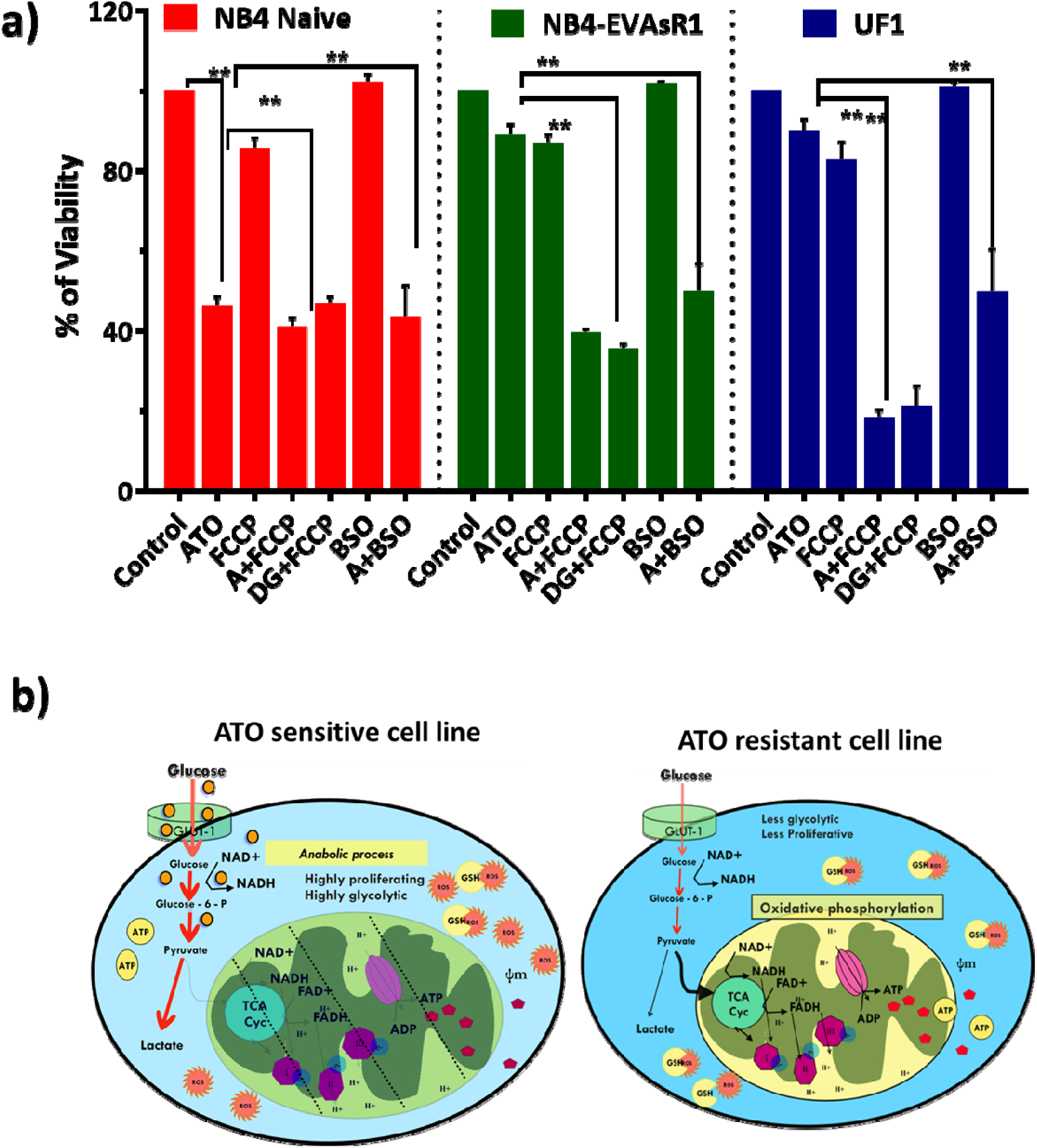
Mitocans synergize with ATO/ 2-DG to promote apoptosis in the ATO resistant cell lines. a) viability of the sensitive and resistant cell lines when treated with OXPHOS uncoupler and GSH synthesis inhibitor. ATO = 2uM; 2-DG= 5mM; FCCP =10uM and BSO = 100uM; All error bars represent the means ± SEM of four independent experiments. *p≤0.05; **p≤0.01; ***p≤0.001; ***p≤0.0001 b) Schematic representation of metabolic adaptation observed in ATO sensitive NB4 naïve and ATO resistant cell lines (NB4 EV-AsR1 and UF1).

Taken together, our data demonstrate that arsenic trioxide resistance is multi-factorial and are not limited to PML or p53 mutations. The generated ATO resistant cell line would be a useful reagent to further evaluate mechanisms of resistance in leukemia. Targeting the metabolic adaptations seen in ATO resistant cell lines has the potential to overcome such resistance. Specifically, our data suggest that co-treatment of ATO and compounds targeting mitochondrial respiration could overcome ATO resistance, and translating this approach to the clinic needs to be explored further.

## Discussion

The combination of ATRA and ATO in the management of APL has improved the survival and treatment outcome of patients by replacing the myelotoxic conventional chemotherapy. However, a proportion of newly diagnosed (10-20%) and relapsed APL (40-50%) patients’ relapse after standard of care. The resistance and therapy failure observed in the clinic against ATO has been focused on the presence / acquisition of *PML* B2 domain mutations ^31,32^. Our recent publication on the mutational spectrum of relapsed and newly diagnosed APL patients suggests the importance of additional genetic events (FLT3, KRAS, NRAS, ARID1B, p53, and WT1) during disease recurrence; however, it is also noted that mutations resulting in primary or secondary ATO resistance were extremely rare and could not explain the majority of disease relapses after treatment with ATO based regimens^26^. Other mechanisms such as bone marrow microenvironment mediated drug resistance, up-regulation of anti-apoptotic factors, modulation of cellular energy metabolism, and oxidative stress can contribute to therapy resistance.

In comparison to the exisiting ATO resistant cell lines^33,34^ our inhouse generated ATO resistant cell lines are stable, well characterized at the genomic levels and also posses the well known ATO resistance conferring mutation A216V in the B2 domain of the PML-RARA oncoprotein.

In comparison to the naïve NB4, in house generated ATO resistant cell lines overexpressed ATO efflux transporters such as AQP9, MRP4, and ABCA7, which correlated with their inability to accumulate intracellular ATO. We did not observe this phenomenon in primary blasts from relapsed APL patients previously treated with ATO in comparison to newly diagnosed^35^.

We noted the presence of a genetic mutation in the B2 domain of the *PML* gene in our in-house generated ATO resistant cell line that has been reported to confer resistance. It is important to note that these mutations are acquired post ATO treatment, and none of the APL cell lines or relapsed APL patients prior to ATO therapy had the B2 domain mutation^36^. ATO sensitivity has also been reported to be impacted by *p53*, which regulates the nuclear body formation and reactive oxygen species levels. We evaluated the p53 mutation and observed that the p53 mutation was present in the ATO resistant cell lines. Based on our data demonstrating nuclear body formation and the lower reactive oxygen species in ATO resistant cell lines the resistant to ATO is likely to be independent of p53 status in these cell lines.

We observed that the cellular redox system of the resistant cell lines is significantly altered with these cell lines having lower reactive oxygen species, lower proliferative rate, and an increased antioxidant system favoring quiescence and stemness like properties^37^.

ATO sensitive cell line NB4 naive was observed to be more reliant on the Warburg effect for their survival and proliferation, and it was significantly affected by a glycolytic inhibitor (2-DG). We also noted that in the in vivo APL model, 2-DG significantly reduced the leukemic burden comparable to the standard of care (ATO). This further supports that the naïve ATO sensitive APL cells are affected by glycolytic inhibition in vivo validating our in-vitro findings.

The ATO resistant cell lines were observed to be dependent on OXPHOS for their survival and also could switch between two metabolic pathways when one is inhibited. A combination of ATO and mitocans significantly affected the survival of resistant cell lines. This combination of ATO and mitocans was also effective in non-APL AML cell lines in-vitro (data not shown).

The observed metabolic plasticity of the ATO resistant cell lines is similar to the previously reported metabolic phenotype of non-APL AML (non-M3) cell lines^38^. It has been noted that AML cells, in contrast to most cancers, utilize OXPHOS preferentially rather than glycolysis for their survival. Based on the data in the existing literature in AML-non M3 and our own data with these ATO resistant APL cell lines suggests that a combination of ATO and mitocans has the potential to be an effective therapeutic combination in the treatment of both ATO resistant APL and non-M3 AML.

## Acknowledgments

This study is supported by a Wellcome-DBT India Alliance research grant (IA/S/11/2500267) and DBT-COE grant (BT/COE/34/SP13432/2015), New Delhi, India. -VM is supported by the senior fellowship program of Wellcome-DBT India Alliance IA/S/11/2500267 and IA/CPHS/18/1/503930, New Delhi, India. SG, HKP, SD were supported by a senior research fellowship from the Council for Scientific and Industrial Research (CSIR), New Delhi, India. - We acknowledge Intas Pharmaceutical Ltd, India, and NATCO pharmaceutical Ltd, India, for kindly providing us API of pharmaceutical drugs for this study. We acknowledge Prof. Mitradas Panicker, National Center for Biological Sciences, Bengaluru, and Prof. Tamil Selvan, Department of Biotechnology, Anna University, Chennai, for providing access to the seahorse extracellular flux analyzer.

## Author contribution

NB: performed research, involved in designing the study, performed molecular tests, and analyzed data and written the paper

SG: performed research, performed molecular tests, and analyzed data.

EC: performed research, performed molecular tests, and analyzed data.

HKP: performed research, performed molecular tests, and analyzed data

AV: performed research, performed molecular tests, and analyzed data

AAA: performed research, performed molecular tests, and analyzed data

SD: performed research, performed molecular tests, and analyzed data.

SK: performed research and analyzed data.

AK: performed research and analyzed data

NBJ: performed research, performed molecular tests, and analyzed data

PB: performed research and analyzed data.

VM: performed research, designed study, analyzed data, and written the paper.

